# New iPSC resource with long-read whole genome sequencing characterizations for enhanced in vitro modeling

**DOI:** 10.1101/2025.06.17.660113

**Authors:** Laura Scheinfeldt, Anthony Pompetti, Gennaro Calendo, Tatyana Pozner, Christine Grandizio, Gretchen Smith, Kelly Hodges, Neda Gharani, Dara Kusic, Matthew Mitchell, Nahid Turan

## Abstract

Here we present a new iPSC resource of apparently healthy subject biospecimens available to the research community through the National Institute of General Medical Sciences Human Genetic Cell Repository (NIGMS Repository). This resource includes five iPSCs and matched parental cell lines with accompanying publicly available, HiFi whole-genome sequencing data. Structural variant (SV) and single nucleotide variant (SNV) concordance between iPSC and parental lines was generally high; however, we found a notable reduction in concordance between the iPSC reprogrammed with retroviral reprogramming and its parental line consistent with previous work showing newer Sendai approaches to be more robust in preserving genomic integrity. This iPSC resource additionally includes pharmacogenomic and human leukocyte antigen (HLA) gene annotations as well as a set of user-friendly, web-based search tools to visualize and explore SVs and SNVs. This new resource is designed to offer a highly characterized set of *in vitro* models for research into cell-type specific functional characterization of genetic, genomic and pharmacogenomic variation. More generally, these renewable biospecimens and genomic data search tools are available to the scientific community to support high-quality and reproducible biomedical research.

## Introduction

Since their discovery in 2007, human induced pluripotent stem cells (iPSCs) have been an important tool for basic, translational, and clinical research (1). Much like human embryonic stem cells (ESCs), iPSCs exhibit distinctive characteristics such as unlimited self-renewal and pluripotency. Notably, the creation of iPSCs does not involve embryo destruction (2), and iPSCs can be reprogrammed from many cell types, including peripheral blood mononuclear cells (PBMCs) isolated from whole blood, fibroblast cell lines (FCLs), and lymphoblastoid cell lines (LCLs) (3).

iPSCs can be differentiated into a wide range of cell types, including cardiomyocytes, neurons, hepatocytes, and immune cells, creating unprecedented opportunities to study the functional consequences of genetic and genomic variation in disease-relevant cell models (4-8). Until recently, cell models have been characterized for genetic and genomic variation (9) and genomic stability with genome-wide arrays (10-14) and short read whole exome and whole genome sequencing (WGS) (15, 16). However, new long-read sequencing technologies improve genetic and genomic characterization with more comprehensive read alignments, more robust structural variant (SV) characterization (17), and high confidence variant phasing relative to older approaches, offering a new way to enhance *in vitro* cell models for disease studies.

To this end, we present a new human iPSC resource of apparently healthy subject biospecimens available for research through the National Institute of General Medical Sciences (NIGMS) Human Genetic Cell Repository (NIGMS Repository) (https://www.coriell.org/1/NIGMS). The NIGMS Repository, established by NIGMS in 1972 at the Coriell Institute for Medical Research, is a collection of high-quality, renewable, and well-characterized biospecimens available to the biomedical research community (3, 18). This new resource includes iPSCs (n=5) and matched parental cell lines (n=5) all with accompanying publicly available, long-read HiFi WGS data to characterize genetic variation, structural variation, pharmacogenetic and HLA annotations as well as to assess genomic stability and variation concordance between parental cell lines and reprogrammed iPSCs.

This new iPSC resource additionally includes a public set of user-friendly, interactive, web-based search tools to visualize and explore SVs and single nucleotide variants (SNVs) as well as pharmacogenomic and human leukocyte antigen (HLA) gene annotations across the sample collection. More broadly, this novel iPSC resource offers a set of highly characterized cell models for research that benefits from cell-type specific functional and mechanistic characterization of genetic, genomic and pharmacogenomic variation.

## Materials and Methodologies

### Samples

We chose ten NIGMS Repository cell lines from four Personal Genome Project (PGP) participant donors for inclusion in the current study (**Table 1**). Notably, PGP participants consented for public sharing of their genomic data. The sample set includes four lymphoblastoid cell lines (LCLs), one fibroblast cell line (FCL), and five iPSCs. GM23248 (FCL), GM20431(LCL), and GM23338 (iPSC reprogrammed from GM23248) are from the same PGP participant hu43860C; GM24385 (LCL), GM26105 (iPSC reprogrammed from GM24385), and GM27730 (iPSC reprogrammed from PBMCs) are from the same PGP participant huAA53E0 (also known as HG002 in the NIST Genome in a Bottle dataset); GM24143 (LCL) and GM26077 (iPSC reprogrammed from GM24143) are from the same PGP participant hu8E87A9 (also known as HG004 in the NIST Genome in a Bottle dataset); and GM24631 (LCL) and GM26107 (reprogrammed from GM24631) are from the same PGP participant hu91BD69 (also known as HG005 in the NIST Genome in a Bottle dataset). The high molecular weight DNA extracted from each of these cell lines are designated with the HM prefix: HM20431, HM23248, HM23338, HM24385, HM26105, HM27730, HM24143, HM26077, HM24631, HM26107 (**Table 1, Table S1**).

**Table 1.** Sample and data information.

| Subject ID<br>(Personal Genome Project ID) | NIGMS Repository cell line ID | NIGMS Repository HMW DNA ID | Sample information | Total HiFi bases | Total HiFi reads | Average mean read length | Average coverage (x) |
| --- | --- | --- | --- | --- | --- | --- | --- |
| hu43860C | GM20431 | HM20431 | LCL | 57,682,198,252 | 3,094,881 | 18,637 | 18.6 |
| hu43860C | GM23248 | HM23248 | FCL | 51,573,153,708 | 2,813,178 | 18,336 | 16.6 |
| hu43860C | GM23338 | HM23338 | iPSC reprogrammed (retroviral) from GM23248 | 54,268,319,594 | 2,979,457 | 18,215 | 17.5 |
| huAA53E0 | GM24385 | HM24385 | LCL | 51,969,370,055 | 3,002,441 | 17,324 | 16.8 |
| huAA53E0 | GM26105 | HM26105 | iPSC reprogrammed (episomal) from GM24385 | 47,024,165,314 | 2,458,250 | 19,128 | 15.2 |
| huAA53E0 | GM27730 | HM27730 | iPSC reprogrammed (Sendai) from PBMCs | 51,004,032,446 | 2,963,620 | 17,170 | 16.5 |
| hu8E87A9 | GM24143 | HM24143 | LCL | 50,304,085,329 | 2,916,498 | 17,248 | 16.2 |
| hu8E87A9 | GM26077 | HM26077 | iPSC reprogrammed (episomal) GM24143 | 54,090,153,442 | 3,175,171 | 17,036 | 17.4 |
| hu91BD69 | GM24631 | HM24631 | LCL | 53,415,212,445 | 2,794,818 | 19,130 | 17.2 |
| hu91BD69 | GM26107 | HM26107 | iPSC reprogrammed (episomal) from GM24631 | 45,933,561,058 | 2,518,745 | 18,125 | 14.8 |

### Cell culture

#### LCLs

Peripheral blood mononuclear cells (PBMCs) were isolated from whole blood, followed by transformation with Epstein Barr Virus (EBV) to establish independent LCLs. Cultures were maintained at 37°C with 5% CO_2_, with medium changes every 3–4 days. Successfully transformed cells, identified by clumping and proliferation, were expanded into larger vessels, and maintained in growth medium (1640 RPMI, 15% FBS, 1% Glutamax) at 37°C with 5% CO2 and ambient oxygen levels in standard tissue culture flasks. LCLs were grown in the absence of antibiotics and antifungals. Cells were split upon reaching confluence, counted and reseeded at the recommended density. Cell density, viability, and morphology were routinely monitored to ensure consistent growth and the absence of contamination. Specifically, to assess bacterial contamination, cultures were inoculated into blood agar plates as well as Trypticase Soy Broth (TSB) and monitored for turbidity and microbial growth under standard incubation conditions. Fungal contamination was evaluated by inoculation into Sabouraud Dextrose Broth (SDB). Mycoplasma contamination (>90 species) was screened using quantitative real-time PCR (qPCR) with the Applied Biosystems MycoSEQ™ Mycoplasma Detection System (Thermo Fisher Scientific). Cell proliferation and viability were assessed by initial cell counting of the transformed cells and a follow-up count after 4–5 days of culture, enabling calculation of doubling time. DNA identity from all cell lines was confirmed using a set of six microsatellite markers (19). The cells were cryopreserved in freezing medium (1640 RPMI, 30% FBS, 5% DMSO) at 5e+6 cells/cryovial for long-term storage in liquid nitrogen.

#### Fibroblasts

Fibroblasts were derived from skin biopsies processed with the gentleMACS Dissociator (Miltenyi Biotec). Biopsy samples were minced with sterile scalpels and transferred to gentleMACS C Tubes containing 1 mL of Collagenase, followed by incubation at 37°C for 30–35 minutes. The reaction was terminated by adding 1 mL of pre-warmed growth medium (MEM supplemented with 15% FBS, 2 mM L-glutamine, and 1% penicillin/streptomycin). The dissociation process was completed using the gentleMACS Dissociator’s pre-set tissue dissociation protocol. The resulting cell suspension was plated in T25 flasks and cultured at 37°C in a humidified incubator with 5% CO_2_ and 3% O_2_. Medium was replaced every 4–5 days, and cells were subcultured. Following fibroblast generation, the cells were cultured in Eagle’s Minimum Essential Medium (EMEM) with Earle’s salts, non-essential amino acids, 2 mM L-glutamine, and 15% FBS. Cultures were maintained at 37°C with 5% CO2 in standard tissue culture flasks. Upon reaching confluence, cells were detached with trypsin-EDTA, counted, and reseeded at the recommended density. Fibroblast cultures were monitored for contamination using TSB, SDB, and blood agar plates, with mycoplasma detection performed with the Applied Biosystems MycoSEQ™ Mycoplasma Detection System (Thermo Fisher Scientific). Cell proliferation was assessed by a 7-day follow-up count to calculate doubling time, and DNA identity from all cell lines was confirmed using a set of six microsatellite markers (19). For long-term storage, FCLs were cryopreserved in freezing medium (growth medium with 5% DMSO) and stored in liquid nitrogen.

#### iPSC reprogramming, culturing and QC

The lines GM26105, GM26077, and GM26107 were reprogrammed using the episomal method. GM27730 was reprogrammed using the Sendai method, and GM23338 was reprogrammed externally on Matrigel with retroviral vectors containing OCT4, SOX2, KLF4, and cMYC (**Table S1**).

For episomal reprogramming of LCLs with OriP/EBNA1 episomal vectors expressing hOCT3/4 with sh-p53, hSOX2, hKLF4, hL-MYC, LIN28, and EGFP, cells were nucleofected using an Amaxa Nucleofector II device (Lonza) with the U-015 program. The cells were maintained in an incubator at 37°C, 5% CO_2_, and 5% O_2_, with medium changes every other day post-nucleofection. Green fluorescent protein (GFP)-positive cells were monitored to assess nucleofection efficiency. On days 6–7 post-nucleofection, transfected cells were replated on Murine Embryonic Fibroblasts (MEFs)-coated culture dishes. After an additional 1–2 weeks, at least 24 clones were manually selected for expansion. These clones were fed daily until enzymatic passaging and further expansion.

PBMCs were reprogrammed via transduction with four SeV vectors expressing hOCT4, hSOX2, hKLF4, hC-MYC, and EmGFP, using the CytoTune Sendai Reprogramming Kit (Thermo Fisher Scientific). Twenty-four hours after transduction, the medium was refreshed, and cells were cultured for approximately six additional days, with medium changes every other day. GFP-positive cells were examined to estimate transduction efficiency. Cells were ready for harvesting and replating on MEF coated culture dishes around day 3 post-transduction. After an additional 2–3 weeks, colonies reached an appropriate size for transfer, and at least 24 colonies were manually picked and selected for expansion. By approximately passage 10, several cryovials were prepared for cryopreservation.

iPSCs were cultured without antibiotics or antifungals in mTeSR1 medium and maintained at 37°C with 5% CO2 and ambient oxygen levels. Cells were split when reaching ∼75-85% confluency at a ∼1:3-1:6 ratio. Subcultivation involved detaching cells using standard dissociation reagents (Versene or ReLeSR) and reseeding in fresh Matrigel-coated vessels with mTeSR1 medium. Medium was replaced daily to maintain optimal conditions. All procedures were performed under sterile conditions, and cell density and morphology were monitored routinely to ensure healthy proliferation and absence of contamination. Cells were frozen in a medium composed of 90% KnockOut Serum Replacement and 10% DMSO.

To ensure the quality and integrity of iPSCs, comprehensive quality control measures were implemented for each reprogrammed cell line. These included microbial sterility testing via blood agar inoculation and broth cultures to check for contamination, alongside mycoplasma testing. Surface antigen (SSEA4) staining verified the undifferentiated state of iPSCs, while embryoid body (EB) formation and RT-PCR confirmed their differentiation potential and gene expression profiles. Alkaline phosphatase staining further validated the presence of undifferentiated stem cells. In addition, no SeV genome or reprogramming factor integration was detected with qRT-PCR in the cell line reprogrammed using the Sendai method. Absence of episomal reprogramming factor integration was confirmed during passages 15-20 with genomic PCR.

#### HMW DNA extraction and QC

For LCLs, FCLs, and iPSCs we performed high molecular weight (HMW) DNA extraction using the Nanobind CBB Big DNA Kit (Pacific Biosciences) on the KingFisher Flex automation system (Thermo Fisher Scientific). The concentration of double stranded DNA was measured with a Qubit Fluorometer (Thermo Fisher Scientific) using the Qubit dsDNA BR Assay Kit (Thermo Fisher Scientific). The 260/280 ratio was measured by spectrophotometry using a single channel Nanodrop (Thermo Fisher Scientific). In addition, we used pulsed-field gel electrophoresis to estimate the molecular weight of the isolated genomic DNA. Briefly, the following settings on CHEF Mapper XA Pulsed Field Gel Electrophoresis (Bio-Rad) system were applied: voltage gradient -5.5 v/cm; run time - 22 hours; included angle – 120 degrees; initial switch time – 35 seconds; final switch time – 90 seconds; ramping factor- “0” (linear). Samples were analyzed on 1% agarose gel using SYBR gold staining. The results were visualized using iBright Imaging Systems (Invitrogen). DNA identity from all cell lines was confirmed using a set of six microsatellite markers (19).

#### Sequencing

High molecular weight DNA was shipped frozen to the University of Washington Long Reads Sequencing Center for Pacific Biosciences HiFi sequencing, including the SMRTbell library sample preparation, with a target of 16-24X genome coverage.

#### Data processing

We used the Pacific Biosciences snakemake workflow for read alignment, variant calling, and phasing as detailed in their tutorial (https://github.com/PacificBiosciences/pb-human-wgs-workflow-snakemake/blob/main/Tutorial.md). We used NIST Genome in a Bottle (GIB) data for HG002, HG004 and HG005 for benchmarking variant quality (**Table S2**).

We performed our initial data review for the subset of our samples that also had GIB reference data (20). For these comparisons, we focused on comparisons of SNV and SV calls between the respective GIB reference files and our corresponding sample data from the same subjects in each GIB reference genome confident region (20) (https://github.com/genome-in-a-bottle/genome-stratifications).

#### Data analysis

We used the hap.py tool (20) (https://github.com/Illumina/hap.py) to conduct pairwise comparisons of all non-reference heterozygous and homozygous SNVs and small (up to 50 bp) insertion/deletions (indels) with GQ>20 in GIB not in difficult regions BED file for HG38 (GRCh38_notinalldifficultregions.bed) between sample datasets derived from the same subject (**Table 1**). The details on how the GIB BED file was created can be found in the GRCh38_Union_README.md file available at the following website (https://ftp-trace.ncbi.nlm.nih.gov/ReferenceSamples/giab/release/genome-stratifications/v3.5/GRCh38@all/Union/GRCh38_Union_README.md, last accessed 4/8/25). We used the Jasmine tool (21) (https://github.com/mkirsche/Jasmine) to conduct pairwise comparisons of all SVs between sample datasets derived from the same subject (**Table 1**).

#### Data annotation

We used two complementary approaches for pharmacogenetic (PGx) annotation. We annotated all PharmVar pharmacogenes excluding *CYP2D6* using our phased variant call file (VCF) and ursaPGx (22). As input, we used the full genomic variant call format (gVCF) for each sample to ensure that all reference alleles were included. We converted all unphased homozygous genotypes to phased within ursaPGx. We used pb-StarPhase to annotate *CYP2D6* and HLA genes using the phased VCF and binary alignment map (BAM) files (23) (https://github.com/PacificBiosciences/pb-StarPhase).

## Results

We collected HiFi long-read whole genome sequencing data from ten samples (**Table 1**). Total HiFi bases ranged from 45,933,561,058-57,682,198,252, total HiFi reads ranged from 2,458,250-3,175,171, average mean read length ranged from 17,036-19,130, and average coverage ranged from 14.8-18.6x.

### SNV concordance across samples from the same subject

We compared SNVs and small (<50bp) indels across sample datasets derived from the same subject (**Table 1**) for all SNVs present in confident mappable regions with a QC>20 (see Methods Section for additional detail). The total number of SNVs called in each sample ranged from 2,872,997-2,946,936. The proportion of concordant SNVs ranged from 85% to 98%, the proportion of SNVs missing in one of the two samples included in the pairwise comparison ranged from 2% to 15%, and the proportion of discordant SNVs was consistently < 1% (**Table 2, Figure 1**). We note that pairwise SNV comparisons that included HM23338 were consistently less concordant (85%) relative to all other sample comparisons, which ranged from 97%-98%.

**Table 2.** SNP and indel comparisons.

| Comparison | # SNPs | # Concordant SNPs | # Missing SNPs in one sample | # Discrepant SNPs | # indels | # Concordant indels | # Missing indels in one sample | # Discrepant indels |
| --- | --- | --- | --- | --- | --- | --- | --- | --- |
| HM23248-HM20431 | 2888299 | 2830339 | 57313 | 647 | 182424 | 174224 | 8160 | 40 |
| HM24143-HM26077 | 2946936 | 2890690 | 56078 | 168 | 185888 | 177494 | 8382 | 12 |
| HM24385-HM27730 | 2916056 | 2835130 | 80626 | 300 | 184127 | 173951 | 10154 | 22 |
| HM24385-HM26105 | 2914629 | 2835923 | 78373 | 333 | 183911 | 173615 | 10279 | 17 |
| HM24631-HM26107 | 2884083 | 2799297 | 84539 | 247 | 182601 | 171654 | 10932 | 15 |
| HM26105-HM27730 | 2912013 | 2811922 | 99590 | 501 | 183526 | 171407 | 12086 | 33 |
| HM20431-HM23338 | 2879780 | 2434387 | 443650 | 1743 | 181169 | 138720 | 42348 | 101 |
| HM23248-HM23338 | 2872997 | 2430837 | 440840 | 1320 | 180092 | 137639 | 42374 | 79 |

Concordant SNPs and concordant indels represent SNPs and indels that are the same genotype in both samples
Missing SNPs in one sample and missing indels in one sample indicate a non-reference genotype call in one sample and a “nocall” in the other sample
Discrepant SNPs and discrepant indels indicate when the genotype calls in both samples are different (e.g., 0/1 vs. 1/1)

**Figure 1.**
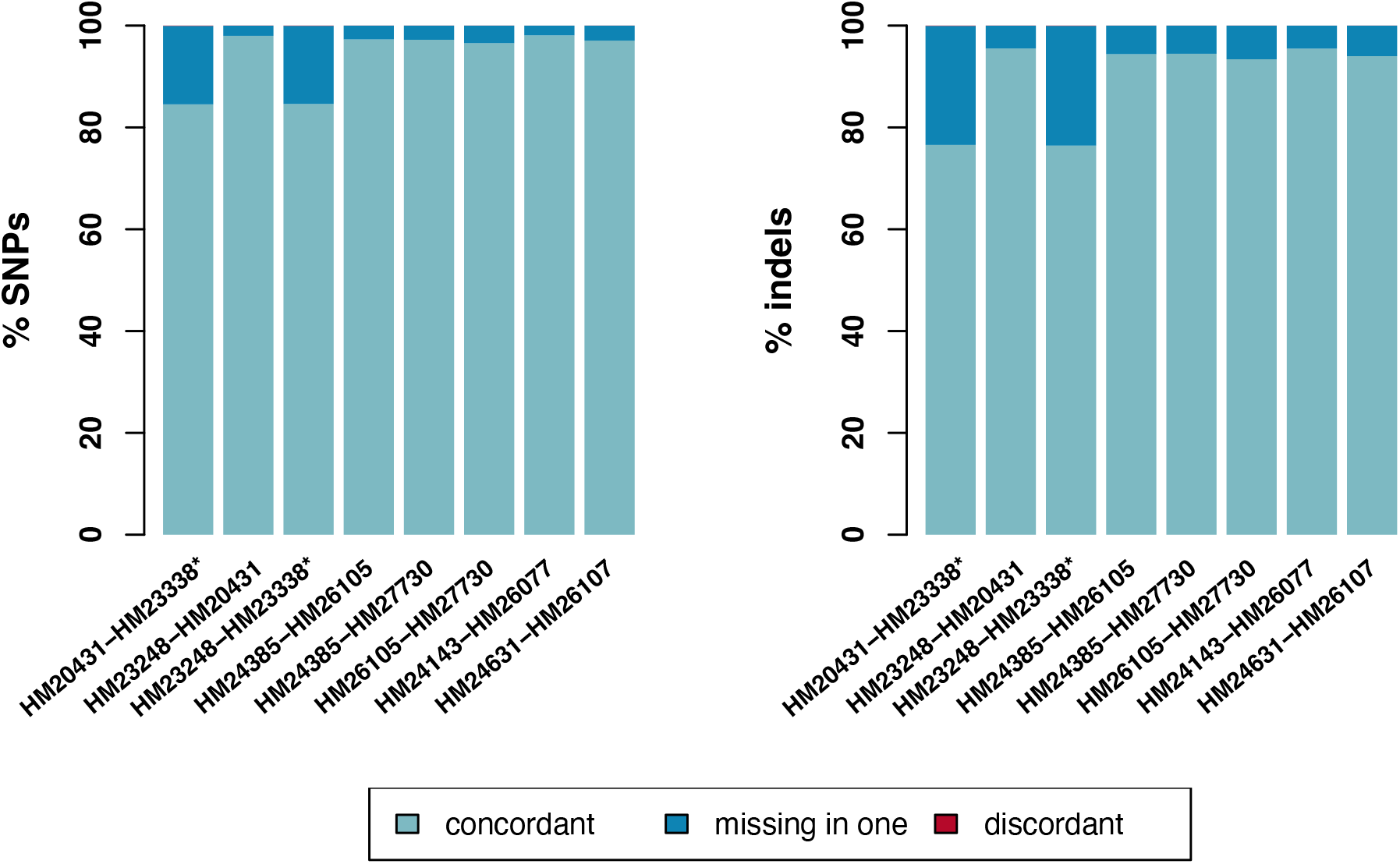
Pairwise SNV/indel comparisons between cell lines from the same subject. The lefthand panel displays pairwise SNV concordance between cell lines from the same subject. HM23338 is denoted with a star because it is the only iPSC in the resource that was reprogrammed externally with retroviral reprogramming. The percent of concordant variants in each pairwise comparison is visualized in light blue, the percent of variants that were missing in one of the two samples included in each pairwise comparison is visualized in blue, and the percent of discordant variants included in each pairwise comparison is visualized in red but is too small to see.

The total number of indels called in each sample ranged from 180,092-185,888. The proportion of concordant indels ranged from 76% to 95%, the proportion of indels missing in one of the two samples included in the pairwise comparison ranged from 5% to 24%, and the proportion of discordant indels was consistently < 1% (**Table 2, Figure 1**). We note that pairwise indel comparisons that included HM23338 were consistently less concordant (76-77%) relative to all other sample comparisons, which ranged from 93%-96%.

To assess the distribution of non-concordant SNVs and small indels across the genome, we performed a sliding window analysis of non-overlapping 1Mb regions. Non-concordant SNVs and small indels were defined as any variant that was either discordant or missing in one of the two samples included in the pairwise comparison. **Table S3** displays the regions of the genome where we found the highest number (<0.001 percentile of the genome-wide distribution) of non-concordant pairwise comparisons of samples from the same subject for SNVs and indels; 76% of top percentile SNV non-concordant regions and 100% of top percentile indel non-concordant regions included a pairwise comparison with HM23338.

### SV concordance across samples from the same subject

The total number of SVs called in each sample ranged from 54,013-56,038, with ranges for duplications, deletions, inversions and insertions ranging from 3,158-3,242, 23,011-24,025, 98-116, and 27,618-23,950, respectively (**Table 3**). Concordance for duplications, deletions, inversions and insertions ranged from 62-78%, 71-89%, 59-76%, and 67-84%, respectively (**Figure 2**). We note that pairwise comparisons that included HM23338 were consistently less concordant (62-77%) relative to all other sample comparisons, which ranged from 73-78%, 83-89%, 74-76%, and 80-84%, respectively for duplications, deletions, inversions and insertions.

**Table 3.** SV comparisons.

| Comparison | total #SVs | # Duplications | # Concordant duplications | # Deletions | # Concordant deletions | # Inversions | # Concordant inversions | # Insertions | # Concordant insertions |
| --- | --- | --- | --- | --- | --- | --- | --- | --- | --- |
| HM23248-HM20431 | 54647 | 3156 | 2314 | 23578 | 19504 | 104 | 77 | 27663 | 22030 |
| HM23338-HM20431 | 54013 | 3158 | 1972 | 23011 | 16442 | 98 | 58 | 27618 | 18366 |
| HM23338-HM23248 | 54351 | 3208 | 2096 | 23189 | 17778 | 101 | 62 | 27743 | 19638 |
| HM26105-HM24385 | 55009 | 3172 | 2483 | 23717 | 21026 | 103 | 83 | 27881 | 23459 |
| HM27730-HM24385 | 54897 | 3158 | 2478 | 23637 | 21033 | 104 | 79 | 27846 | 23430 |
| HM27730-HM26105 | 54863 | 3163 | 2466 | 23549 | 20692 | 104 | 83 | 27923 | 23115 |
| HM24143-HM26077 | 56038 | 3242 | 2534 | 24025 | 21356 | 116 | 88 | 28527 | 23950 |
| HM24631-HM26107 | 54998 | 3143 | 2447 | 23517 | 20713 | 105 | 78 | 28091 | 23474 |

**Figure 2.**
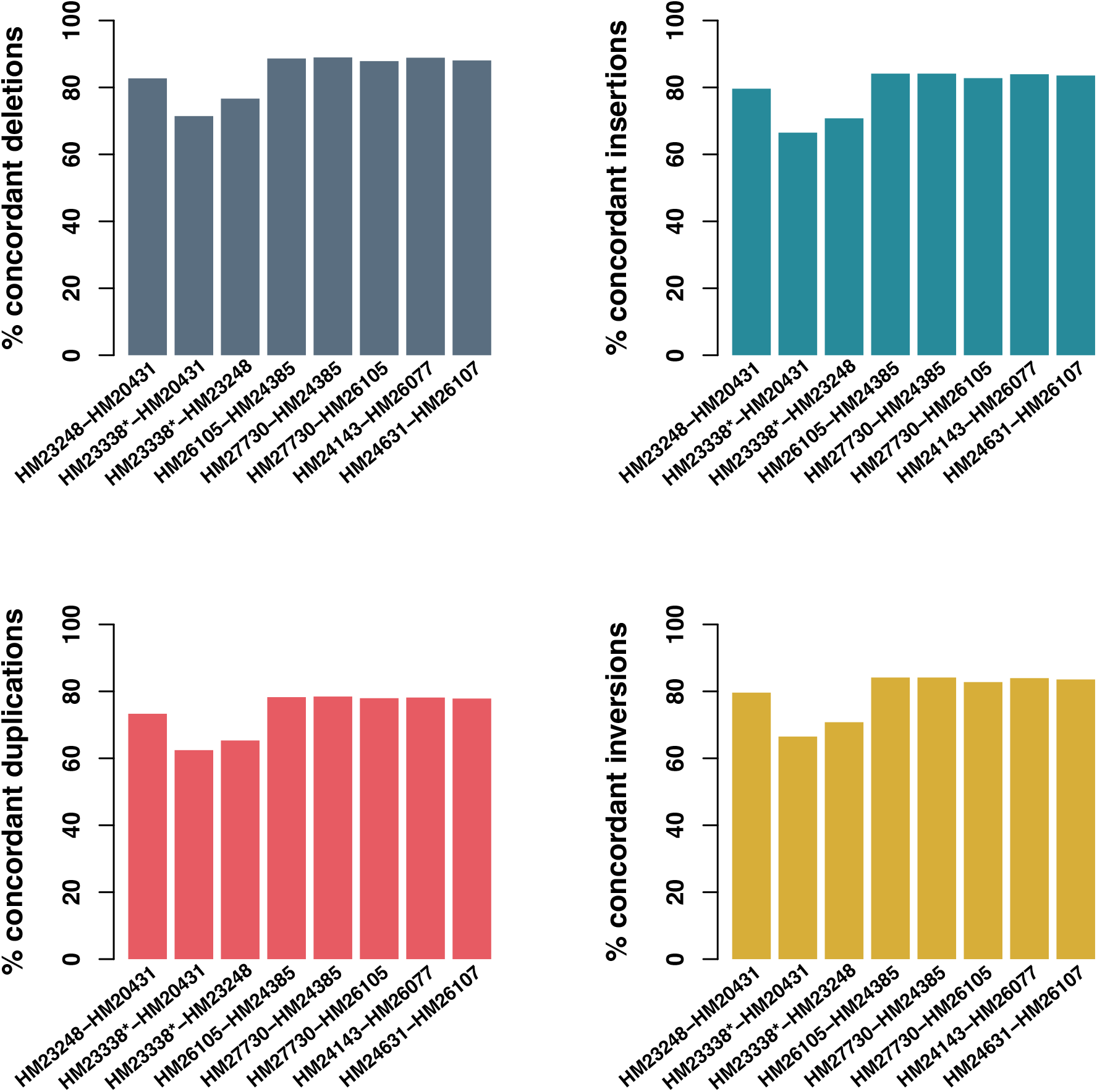
Pairwise structural variation comparisons between cell lines from the same subject. The lefthand top panel displays pairwise deletion concordance in dark blue, the righthand top panel displays pairwise insertion concordance in blue, the lefthand bottom panel displays duplication concordance in red, and the righthand bottom panel displays inversion concordance in yellow between cell lines from the same subject. HM23338 is denoted with a star because it is the only iPSC in the resource that was reprogrammed externally with retroviral reprogramming.

We defined non-concordance as any SV that was present in only one of the two samples included in the pairwise analysis. To assess how non-concordant SVs were distributed across the genome, we performed a sliding window analysis of non-overlapping 1Mb regions. **Table S4** displays the regions of the genome where we found the highest number (<0.001 percentile of the genome-wide distribution) of non-concordant SVs in pairwise comparisons of samples from the same subject for each SV category (deletions, duplications, insertions, inversions). We note that 44% of deletion, 29% of inversion, 40% of duplication, and 42% of insertion non-concordant regions, respectively involve a pairwise comparison with HM23338.

Pharmacogenetic and HLA annotation

Table 4 displays the pharmacogenetic (PGx) and HLA annotations. All overlapping pharmacogene (*CYP2B6, CYP2C19, CYP2C9, CYP3A5, CYP4F2, DPYD, NUDT15, SLCO1B1*) annotations were concordant with those previously reported for HG002, HG004, and HG005 by Holt et al. (23) for all comparisons excluding HG005 *NUDT15* star allele annotations. *NUDT15* exon 1 includes a 6-bp (GAGTCG) repeat that is present with three repeats on the *1 haplotype, four repeats on the *2 and *6 haplotypes, and two repeats on the *9 haplotypes (24). Given this complexity, we manually reviewed the sequencing BAM files for each sample to confirm the number of 6-bp repeats. HM24631 and HM26107 both have four 6-bp repeats on both alleles, and rs116855232 on one allele; we therefore corrected the ursaPGx *NUDT15* annotation output from *1/*3 to *2/*6. All other samples in the collection were manually confirmed to have three 6-bp repeats. We note that one set of subject samples (HM20431, HM23248, HM23338) has a single ambiguous *CYP2A6* allele in the output from ursaPGx, and this is due to the presence of a combination of variants that do not exactly match any PharmVar annotated star allele (both rs1809810 and rs56314118 non-reference alleles on the same phased haplotype), consistent with the approach of ursaPGx as previously described (22); however, this haplotype does match the PharVar *CYP2A6*18*.*002* suballele, and we manually annotated these sample suballeles in **Table 4**. More generally, we note that all PGx diplotype annotations among samples from the same subject annotated with ursaPGx were concordant (the ‘haplotag’ assignment of haplotypes is arbitrary in the absence of the maternal and paternal parental haplotype information, which we did not include in our analysis).

**Table 4.**
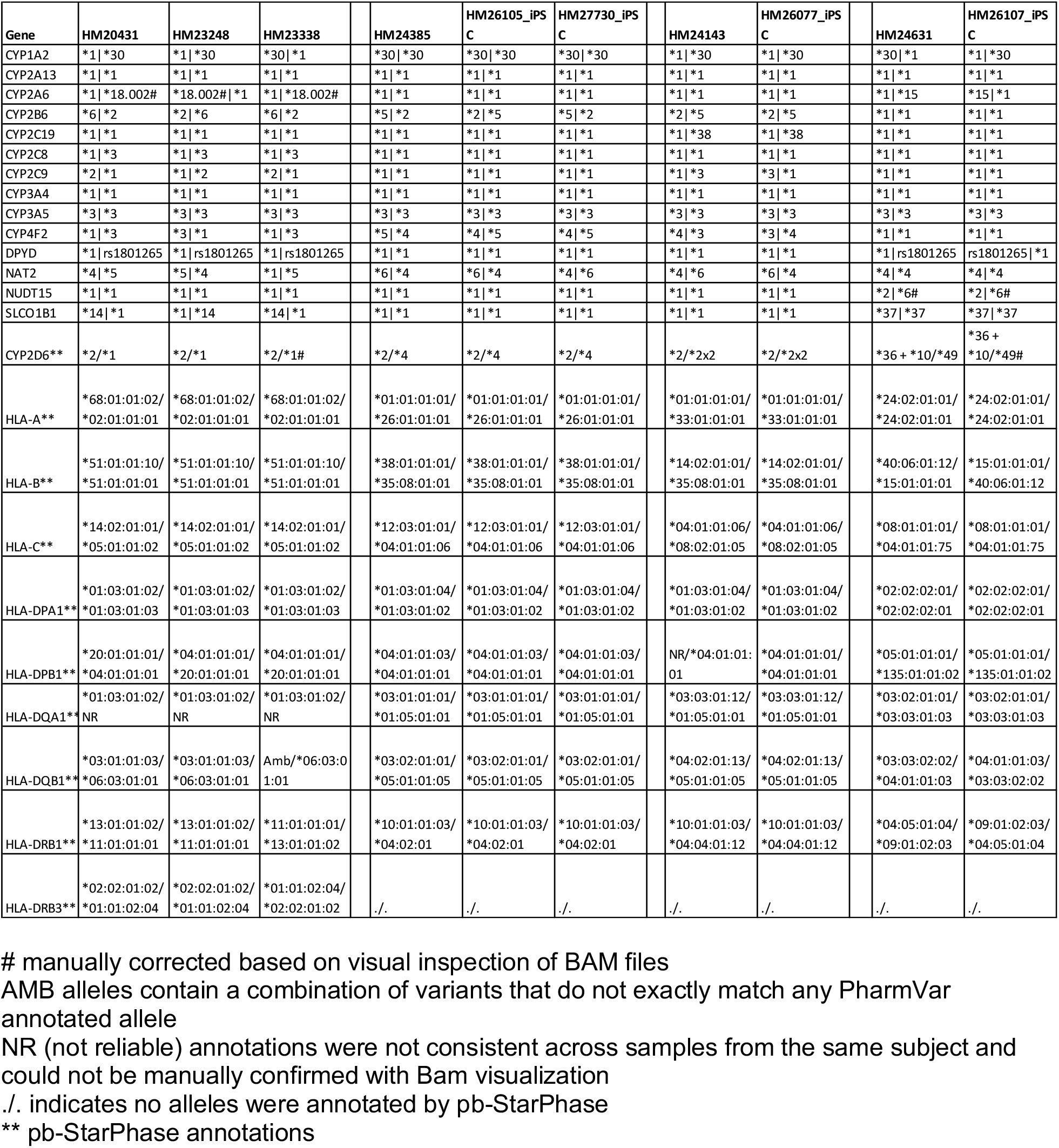
Consensus PGx and HLA annotations.

*CYP2D6* and HLA gene annotations were performed with pb-StarPhase (23) using both the phased VCF and BAM files. The vast majority (>99%) of annotations among samples from the same subject were concordant (**Table 4**). In addition, annotations of samples from Subject HG002/NA24385/GM24385 share at least one allele with HG004/NA24143/GM24143 annotations, consistent with their parent/child relationship. However, we found two *CYP2D6* discrepancies across samples from the same subject: HM23338 has three copies of the *CYP2D6*\*2 allele (defined by rs1135840 and rs16947) relative to HM23248 and HM20431, which were both annotated as *1/*2; HM26107 has one additional variant copy (rs1135822) that defines the *CYP2D6*\*49 allele (relative to *CYP2D6*\*10) that is not present in HM24631. Manual inspection of the BAM file reads supports the HM23248, HM20431 and HM24631 *CYP2D6* annotations; the HM23338 and HM26107 *CYP2D6* annotation inferences are based on lower than average coverage at their respective star allele defining variants (six reads for rs1135840 and 11 reads for rs16947, respectively in HM23338; 12 reads for rs1135822 in HM26107). We additionally found three HLA gene discrepancies across samples from the same subject: the *HLA-DQA1*\*05 allele is not consistent across HM20431/HM23248/HM23338; the *HLA-DQB1*\*03 allele is not consistent in HM23338; and *HLA-DPB1* is not consistent between HM24143/HM26077 for one allele. *HLA-DRB4* and *HLA-DRB5* are not included in **Table 4** because there was no complete set of annotated alleles for either gene for any subject sample set.

### Web based search tools

As part of this new public resource of NIGMS Repository iPSCs, we have created a set of user friendly, web-based search tools of single nucleotide and insertion/deletion variation, and structural variation found in each cell line (https://genomicviewer.coriell.org/). These tools utilize a JBrowse framework (25, 26) that facilitates searches of sample BAM, SNV VCF and SV VCF files by rsid, gene name, or by chr and position range and visualizes the variants found in the dataset in an interactive genomic viewer interface. Users can choose all or a subset of samples to visualize, can choose whether to include only BAM files, only VCF files or both, and can zoom in or out of any region of interest (**Figure 3**).

**Figure 3.**
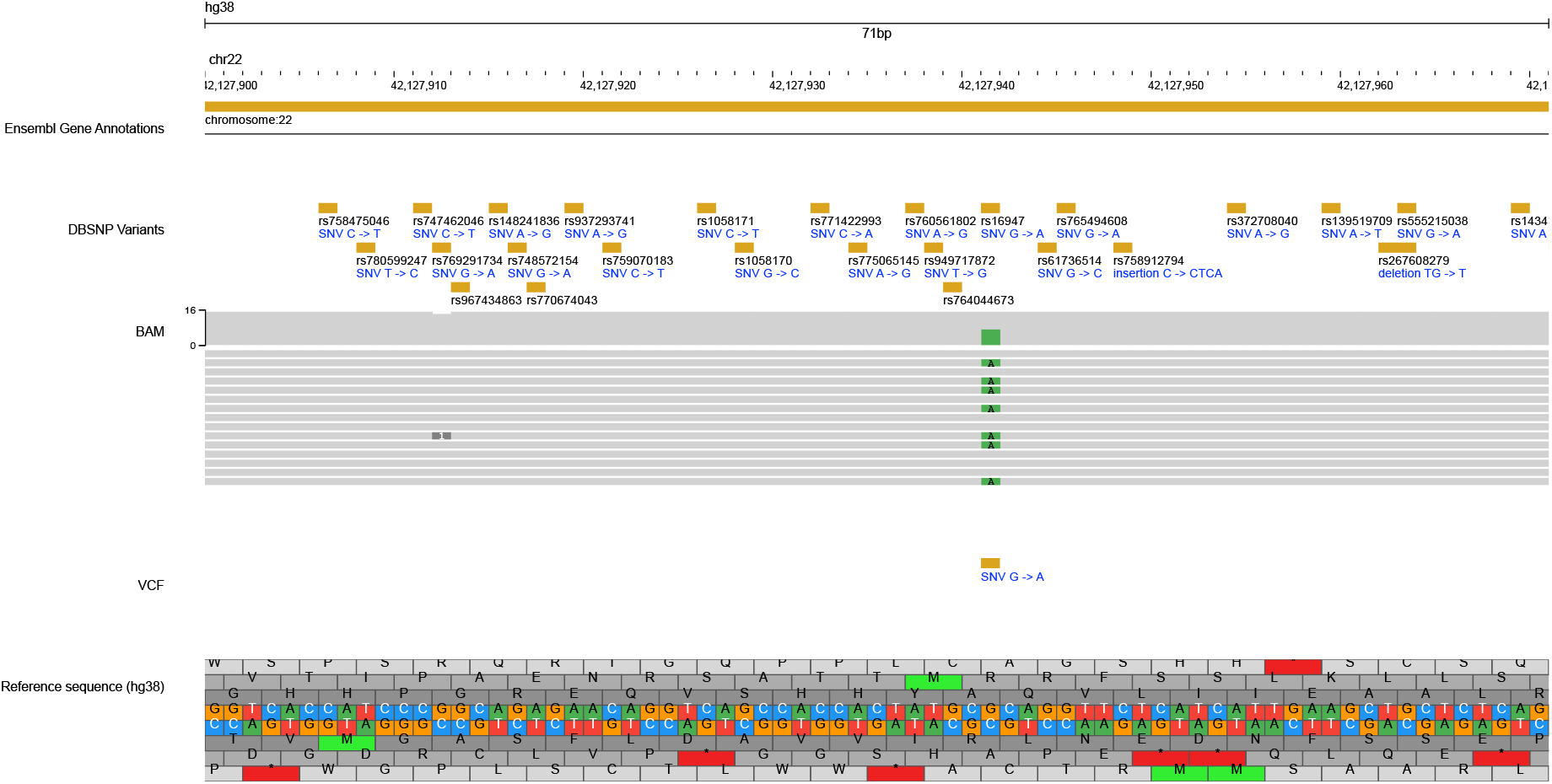
Example iPSC WGS Search Screenshot. Figure 3 displays an iPSC WGS Search example of a 70 base pair region of chromosome 22 (chr22:42,127,900..42,127,970). This search includes the hg38 reference sequence track, the DBSNP track, and the BAM and SNV VCF tracks for HM24385.

## Discussion

Here we describe a new publicly available resource for a collection of NIGMS Repository iPSCs from four subjects and their respective parental cell lines (LCL or FCL). This collection has been characterized with publicly available, long-read whole genome sequencing data as well as HLA and pharmacogenomic annotations. For researchers that prefer not to mine these large-scale datasets directly, this iPSC resource includes a set of web-based search tools with a user friendly, graphical user interface. To our knowledge, this is the only iPSC resource of its kind.

In general, we found high SNV and small (<50 bp) indel concordance between samples from the same subject (**Table 2, Figure 1**); the only pairwise concordance below 97% included HM23338. We note that HM23338 was extracted from GM23338, which is the only iPSC included in the collection that was reprogrammed externally with retroviral reprogramming, which is one of the earlier reprogramming methods that has previously been shown to be vulnerable to insertional mutagenesis and viral integration into the host genome (3, 27-31). These reprogramming methodological vulnerabilities are the most likely explanation for the relatively low HM23338 pairwise concordance (85%).

As expected, SV concordance across all included samples from the same subject (**Table 3, Figure 2**) varied by SV type and was highest for insertions and deletions, relative to duplications and inversions. Similar to the SV results, and presumably for the same reprogramming methodology vulnerabilities (3, 27-31), when excluding comparisons to HM23338, pairwise concordance of all SV types increases (73.3%, 82.7%, 74.0%, and 80.0% for duplications, deletions, inversions, and insertions, respectively). There is an additional, relatively modest increase in SV concordance when further excluding the FCL-derived HM23248 (77.9%, 87.9%, 74.3%, and 82.8% for duplications, deletions, inversions, and insertions, respectively). This increase is potentially due to previously noted somatic mutations in FCLs (32).

The web-based visualization tools included in this iPSC resource offer a user-friendly way for researchers to further explore SNV and SV concordance and read coverage in each iPSC and matched parental line thus allowing in-depth review and visualization of variants of interest. In the current analysis, we have focused our visual variant inspection of BAM files on pharmacogene regions to ensure that the sample-associated PGx annotations are robust, and we were able to manually confirm annotations for fifteen pharmacogenes (**Table 4**) despite the limited 17x average sequencing coverage.

Taken together, the consistent reduction in SNV and SV concordance between iPSC DNA HM23338 and its parental cell line DNA (HM23248) reinforces the advantages of newer, more robust reprogramming methods such as the Sendai viral reprogramming method for iPSC establishment. Unlike integrating viral vectors that may leave genomic footprints or cause genomic instability, efficient Sendai virus-based reprogramming delivers reprogramming factors transiently without altering the host genome, resulting in iPSCs with higher fidelity to donor cell genetic and genomic profiles (3, 33).

This new collection of characterized and annotated iPSCs and parental cell lines offers an unprecedented opportunity to study genetic, genomic and pharmacogenomic variation in two-dimensional disease-relevant cell cultures and three-dimensional disease-relevant organoids. For example, iPSC-derived hepatocyte like cells (HLCs) offer a complementary *in vitro* system to primary liver cells, which are challenging to work with on large-scale drug screening studies (7). Metabolic enzyme expression, such as cytochrome P450 gene expression in iPSC-derived HLCs is lower relative to primary hepatocytes, but more scalable for high-throughput drug screening and liver toxicity studies (7). In addition, ongoing developments in differentiated HLC generation and handling are improving HLC maturation and related gene expression patterning (8). Similar applications involving disease-focused studies on heart disease, psychiatric, neurological, and neurodegenerative disorders, as well as immune, metabolic, and gastrointestinal diseases will likely benefit from apparently healthy control iPSCs with public genetic, genomic, and pharmacogenomic characterizations (4-6). More generally, any research focused on tissue-specific functional characterization of SVs and SNVs will benefit from this new collection that complements and reduces the need for animal models in research and drug screening.

In conclusion we present a novel resource of iPSCs and matched parental cell lines sequenced with publicly available, long-read HiFi WGS with characterized genetic variation, structural variation, and pharmacogenetic and HLA annotations. This resource is accompanied by a set of user-friendly, web-based search tools to enable scientists utilizing the resource to explore and visualize single nucleotide and structural variation. Taken together, this iPSC resource offers a set of highly characterized *in vitro* model systems for cell-type specific functional characterization of genetic, genomic and pharmacogenomic variation.

## Acknowledgements

We are grateful to the Personal Genome Project participants that generously donated the samples that were used to create this new resource. We would like to thank Jozef Madzo for his advice on the genomic data collection, and we acknowledge the critical contributions of Phillip

Hodges for the design, development, implementation and ongoing support of the IT infrastructure needed for the public genomic search tools.

## Data availability

The raw sequencing data generated in this study are available through SRA at the following link: https://www.ncbi.nlm.nih.gov/sra/PRJNA1273174

The web-based access to browse BAM and VCF files can be found at the following link: https://genomicviewer.coriell.org/

